# The Identification of a 1916 Irish Rebel: New Approach for Estimating Relatedness From Low Coverage Homozygous Genomes

**DOI:** 10.1101/076992

**Authors:** Daniel Fernandes, Kendra Sirak, Mario Novak, John Finarelli, John Byrne, Edward Connolly, Jeanette EL Carlsson, Edmondo Ferretti, Ron Pinhasi, Jens Carlsson

**Affiliations:** School of Archaeology, University College Dublin, Belfield, Dublin 4, Republic of Ireland; CIAS, Department of Life Sciences, University of Coimbra, 3000-456 Coimbra, Portugal; Department of Anthropology, Emory University, 201 Dowman Dr., Atlanta, GA 30322, United States of America; Institute for Anthropological Research, Ljudevita Gaja 32, 10000 Zagreb, Croatia; School of Biology and Environment Science, University College Dublin, Belfield, Dublin 4, Republic of Ireland; Earth Institute, University College Dublin, Belfield, Dublin 4, Republic of Ireland; National Forensic Coordination Office, Garda Technical Bureau, Garda Headquarters, Phoenix Park, Dublin 8, Republic of Ireland.; Forensic Science Ireland, Garda Headquarters, Phoenix Park, Dublin 8, Republic of Ireland; Area 52 Research Group, School of Biology and Environment Science, University College Dublin, Dublin 4, Republic of Ireland

## Abstract

Thomas Kent was an Irish rebel who was executed by British forces in the aftermath of the Easter Rising armed insurrection of 1916 and buried in a shallow grave on Cork prison's grounds. In 2015, ninety-nine years after his death, a state funeral was offered to his living family to honor his role in the struggle for Irish independence. However, inaccuracies in record keeping did not allow the bodily remains that supposedly belonged to Kent to be identified with absolute certainty. Using a novel approach based on homozygous single nucleotide polymorphisms, we identified these remains to be those of Kent by comparing his genetic data to that of two known living relatives. As the DNA degradation found on Kent's DNA, characteristic of ancient DNA, rendered traditional methods of relatedness estimation unusable, we forced all loci homozygous, in a process we refer to as “forced homozygote approach”. The results were confirmed using simulated data for different relatedness classes. We argue that this method provides a necessary alternative for relatedness estimations, not only in forensic analysis, but also in ancient DNA studies, where reduced amounts of genetic information can limit the application of traditional methods.

## INTRODUCTION

Estimating the genetic relatedness of modern individuals is routinely achieved by employing the use of microsatellites (synonymous with short tandem repeats (STR)) or other genomic markers that estimate kinship coefficients based on probabilities of identity-by-descent (IBD)^1,2^. These methods, however, cannot be applied to cases where the DNA presents high levels of fragmentation and damage, as is common in ancient DNA (aDNA) research. Upon an organism's death, its genetic material starts to degrade and accumulate damage as repair enzymes no longer maintain the integrity of the molecular structure of DNA^3,4^. Among the many factors that contribute to the rate and severity of this phenomenon are temperature, the acidity of the surrounding depositional environment, ambient level of humidity, and the eventual invasion of environmental microbes into the organism's cells. As a result, DNA fragments extracted from preserved tissue (in most cases bone and teeth) that is recovered from either ancient or semi-ancient (e.g. many forensic cases) human remains, are short in length, ranging from 30 to 70 base pairs. The degradation process has a major impact on the success rates and authenticity of many PCR-based ancient DNA (aDNA) identification techniques^3,4,5,6^, however analysis of these short and damaged DNA molecules was revolutionised with the onset of Next Generation Sequencing (NGS) one decade ago. Next-Generation shotgun sequencing has enabled aDNA studies to progress at a much faster rate than before, and when applied in conjunction with optimised bone tissue isolation, DNA extraction, and sequencing technologies, large amounts of genetic information can be obtained even from samples with poor molecular preservation.

Relatedness estimation is a topic of relevance and interest in both anthropological and forensic studies. Before NGS, PCR-based studies were affected by a limited capacity to authenticate aDNA results and an inability to retrieve the required data from most aDNA samples^7,8,9,10^. However, some methods have been adapted to work specifically with this type of NGS or ancient DNA data; these are present in software such as PLINK2^11^ and NGSrelate^12^. Both software packages utilise Single Nucleotide Polymorphism (SNP) data, shown to work well with maximum likelihood approaches, and rely on genotypes, genotype likelihoods and minor allele frequencies. However, these packages require the input of relatively high amounts of genetic data (large numbers of loci) which is oftentimes challenging and expensive to obtain from ancient skeletal material^1,2,12^. Our method overcomes these challenges by substantially reducing the amount of input data required without sacrificing the confidence of the relatedness estimation. Here, we apply this novel method to identify the century-old skeletal remains of a famous Irish Rebel, Thomas Kent.

Thomas Kent (1865-1916), an Irish rebel native to Castlelyons, grew up in Bawnard House located just outside the town of Fermoy in County Cork, Ireland. A week after the Easter Rising insurrection, in April of 1916, the Royal Irish Constabulary (RIC) raided the family home on 1^st^ May. An RIC officer was shot dead during the raid. Thomas and William Kent were arrested. Following court martial, William was acquitted, but Thomas received a death sentence and was one of 16 men executed by British Forces following the Easter Rising, being executed in the early hours of the 9^th^ of May, 1916 at Cork Detention Barracks and then buried adjacent to where he fell^13^.

The remains of Thomas Kent lay in the Barracks, which subsequently became Cork Prison, until June 2015, when they were exhumed by a team led by the National Monuments Service of the Department of Arts, Heritage and the Gaeltacht. Poorly kept records from the era of Thomas Kent's execution and throughout the intervening 99 years resulted in confusion surrounding his final resting place and uncertainty in the identification of his remains. The presumed identity of the remains was solely based on circumstantial evidence, and though attempted, it was soon determined that traditional DNA analysis was not an option due to the DNA degradation that was expected to be found in his remains due to their ancient/archaeological origin. The National Forensic Coordination Office at the Garda Technical Bureau and Forensic Science Ireland contacted the University College Dublin (UCD) who developed a new DNA identification method, based on optimisation techniques involving the use of the osseous inner ear part of the petrous part of the temporal bone^14^ which has been applied successfully for over ~1000 archaeological samples from temperate regions spanning between 40,000-500 years before present (average endogenous yields range of 50-70% and with an overall success rate of ~80%^15^).

Using low-coverage shotgun sequencing data obtained from a single sequencing run on the Illumina MiSeq platform, we compared modern genetic data from two of Thomas Kent's living relatives to his century-old genetic material in order to identify his remains. Based on the success of the our analytic approach in a low-coverage data scenario, we propose a NGS shotgun SNP-based method for relatedness estimation that uses “forced homozygote” allele data to estimate relationship coefficients and is based upon a symmetrical Rxy estimator algorithm developed by Queller and Goodnight^16^.

Similar to other available software, the approach reported in this study relies on SNP data but requires a substantially lower amount of input data than the methods mentioned previously while not sacrificing any accuracy. This makes it widely applicable budget-efficient forensic applications, as well as to the rapidly-expanding field of ancient DNA studies, where other methods are not an option, because low coverage homozygous data is the norm^10,^^15^. Here we detail the success of our approach in the identification of the historical remains of the Irish revolutionary Thomas Kent.

## RESULTS AND DISCUSSION

### Authentication of Sequencing Data

As expected, DNA preservation differed noticeably between the modern individuals and the supposed archaeological remains of Thomas Kent (hereafter, TK). Because of that, we followed the methodologies used for ancient DNA analysis. For standardization purposes, after separate DNA extractions, which required different protocols due to the use of different biological tissues, we prepared the modern samples for sequencing in exactly the same way as TK. The average sequence read length from TK was predicted to be shorter than his modern relatives (E81 and E82) due to the historic nature of this sample; average fragment length was determined to be 54.01 base pairs (bp), with a wide standard deviation of ± 11.57bp (Table 1). In contrast, the modern relatives' DNA size averaged 64.48bp, with a standard deviation of ± 1.52bp, which is extremely close to the sequencing length used (65bp). During the analysis of the raw sequencing data, the presence of adapters was detected in very few reads for the modern individuals as compared to the ancient sample (38% for E81 and E82, against 72% for TK), further supporting the notion that these endogenous modern DNA fragments were longer than 65bp. This was the expected outcome for modern DNA samples, indicating that these non-damaged sequences were possibly of lengths greater than or close to 65bp. Due to the archaeological nature of Thomas Kent's genetic material and the possibility of modern DNA contamination, raw data for this sample was first analysed to confirm the authenticity of the retrieved DNA as endogenous and ancient. To authenticate the DNA of TK as ancient, we utilised a widely-used approach developed for ancient DNA that quantifies deamination frequencies at the terminal ends of the DNA molecule, looking in particular for C>T substitutions at 5' overhangs that characterize the deamination of cytosines. Using the mapDamage v 2.0 software^17,^^18^, the deamination frequencies present in TK's DNA, 0.14 C>T at the 5' end and 0.10 G>A at the 3' end (Figure 1) appear consistent with the expectation of molecular degradation for century-old bones interred in a shallow grave in the presence of quicklime^13^. In contrast, the modern DNA from TK's living relatives did not show significant damage patterns at the ends of the sequences. However, because fragments with shorter size than that of the sequencing length (65bp) are not expected to be overwhelmingly present in modern DNA, these deamination frequencies are not informative as it is probable that the ends of the molecules were not read.

**Figure 1:**
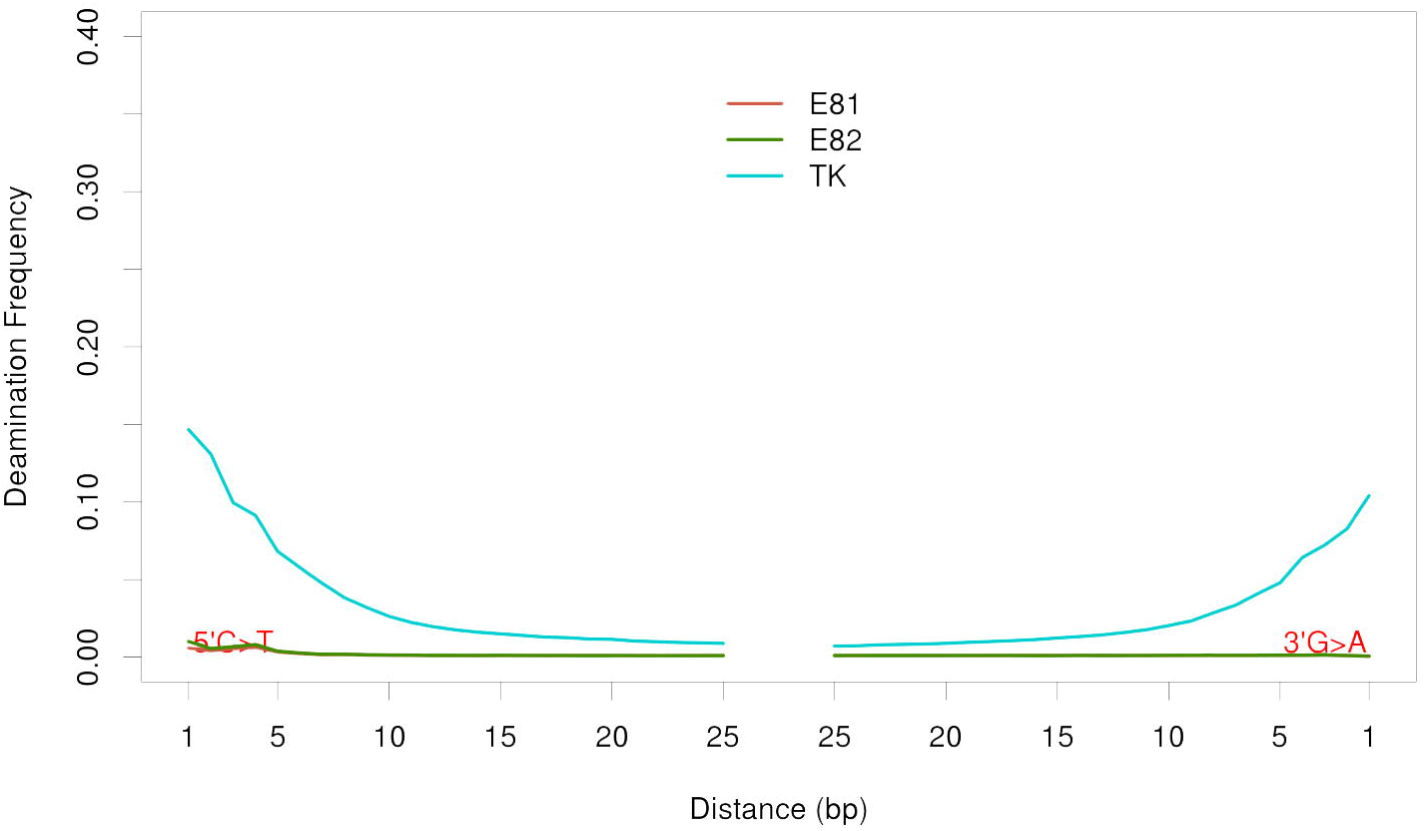
DNA damage patterns from deamination frequencies of terminal bases.

**Table 1:**
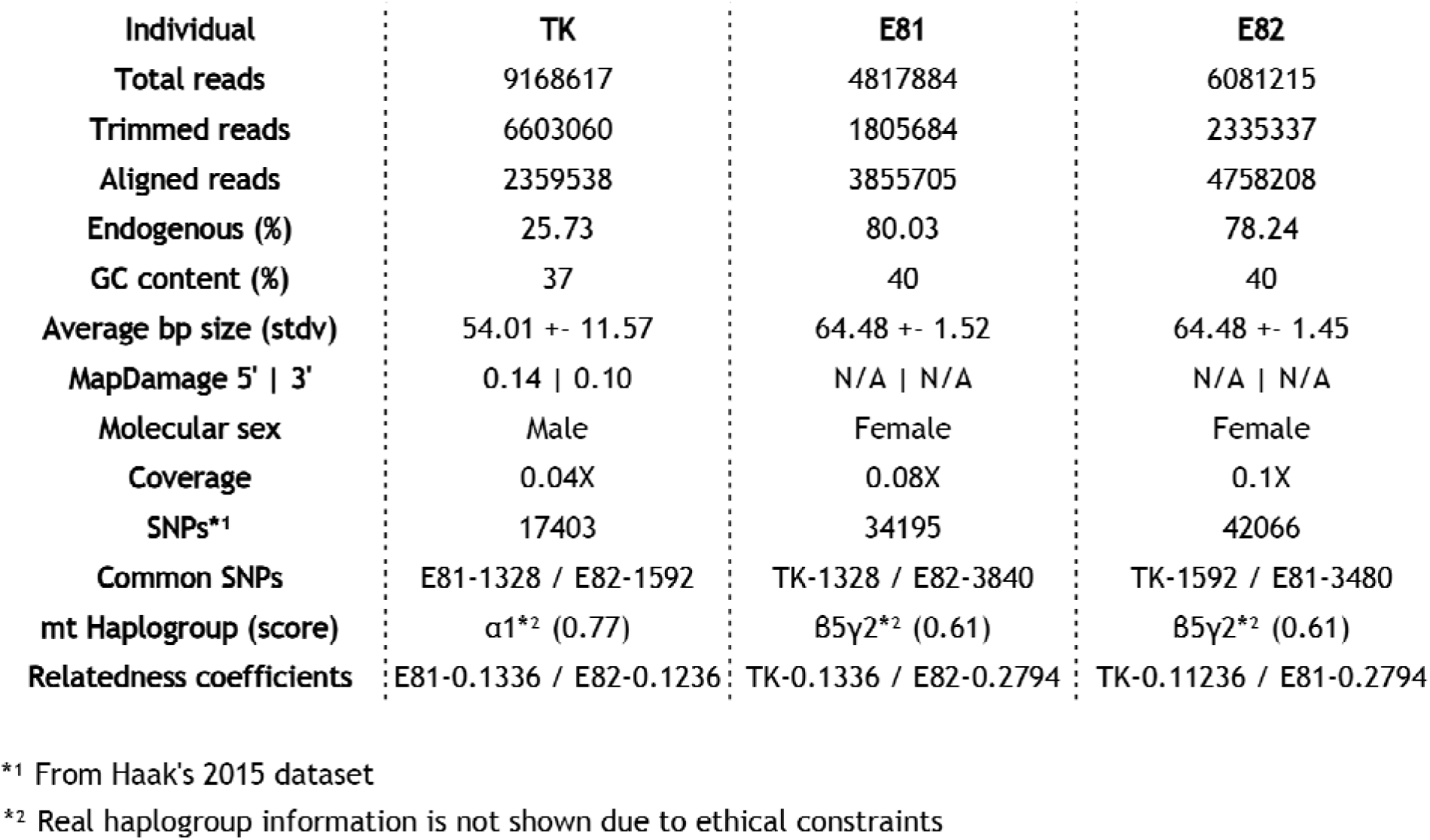
Sequencing data analysis and relatedness coefficient results.

Mitochondrial haplogroups were also estimated for the three individuals (Table 1) using Phy-Mer^19^. For ethical reasons, the determined haplogroups are not reported in this paper; however, the two modern relatives, as expected, shared the same haplogroup, with a score of 0.61. This score estimates how well the given data matches the assigned haplogroup in the 0-1 interval. Thomas Kent's haplogroup was different from that of the relatives, with a score of 0.77. All three haplogroups were consistent with expectations for historic or modern individuals native to Ireland.

### Relatedness Estimations

We estimated relatedness among the TK remains and two surviving members of the Kent family using very low coverage shotgun data (ranging from 0.04X to 0.1X) obtained from one MiSeq sequencing run, which currently generates a maximum of 25 million reads. Because we did not use a targeted enrichment or hybridization capture method to selectively identify and obtain common loci within the human genome, the output data for each individual was a random pool of overlapping reads. Along with the negative controls, these three samples were the only samples placed on the sequencing run. Thomas Kent had 25% of his total reads aligning to the human genome, representing a genomic coverage of 0.04X.

This amount of endogenous DNA is considered high in an ancient DNA context and was made possible to retrieve because of improved DNA extraction methodologies^14, 20^. Relative 1 had 4817884 total reads, with 3855705 aligning to the human genome (80% endogenous contents and 0.08X coverage) and Relative 2 slightly more, 6081215 total reads, from which 4758208 were of human origin (78% endogenous contents and 0.1X coverage) (Table 1). None of the negative controls prepared along with the samples rendered human sequences. Using the dataset of SNPs developed for population and evolutionary genetic studies employed in^10^, we called genotypes for 354,212 positions for each individual, obtaining 17403 SNPs called for TK, 34195 for Relative 1 (E81), and 42066 for Relative 2 (E82). Out of these, we extracted only the shared SNPs between each dyad: TK:E81 (1328 SNPs), TK:E82 (1592 SNPs), and E81:E82 (3480 SNPs). As the total genome coverages were very low (Table 1), virtually all SNPs called had only one 1X read depth. Because we did not have more than one read per SNP position, we forced each SNP to be homozygous by repeating the called base to generate a diploid loci; this is referred to as the “forced homozygote” approach. For SNPs with more than 1X coverage, one call with phred quality above 30 was randomly selected and then “forced” homozygous by repeating the base as explained above. We estimated relationship coefficients for each of the three dyads using the Queller and Goodnight (1989) algorithm incorporated in the software SPAGeDi1-5a (build04-03-2015)^21^, using the correspondent European allele frequencies downloaded from the 1000 Genomes website. As anticipated, the use of the forced homozygote approach resulted in relatedness coefficients (Rxy) of half the expected values. For the pair E81:E82, we observed an Rxy of 0.2794, consistent in modern genetics with second order relatedness, but with first order relatedness in the forced homozygote approach (i.e. equivalent to a modern genetics Rxy of 0.50). For TK:E81, the Rxy was estimated at 0.1336, and for TK:E82, it was estimated at 0.1236. These values are consistent with a second order of relatedness for uncle/niece (25% in modern genetics, or 12.5% under our forced homozygote approach) between TK and the two living relatives, supporting the positive identification of his remains.

The expected hypothesis that Thomas Kent was related to the two living relatives by a second order relationship and the two living relatives are related to each other by a first order relationship (Hypothesis #4, Table 2) is unambiguously supported by the data, comprising nearly the entire posterior probability of the set of hypotheses. Using the posterior probabilities, the odds that this hypothesis is incorrect given the observed data is less than one in one million (8.15 E-07). Indeed, the Odds Ratio of the summed posterior probabilities for the four hypotheses proposing that the remains of Thomas Kent are related, in any manner, to both relatives versus the odds that he is unrelated to at least one of the two is in excess of 5 trillion, indicating conclusively that the TK remains are related to the two living members of the Kent family.

**Table 2:**
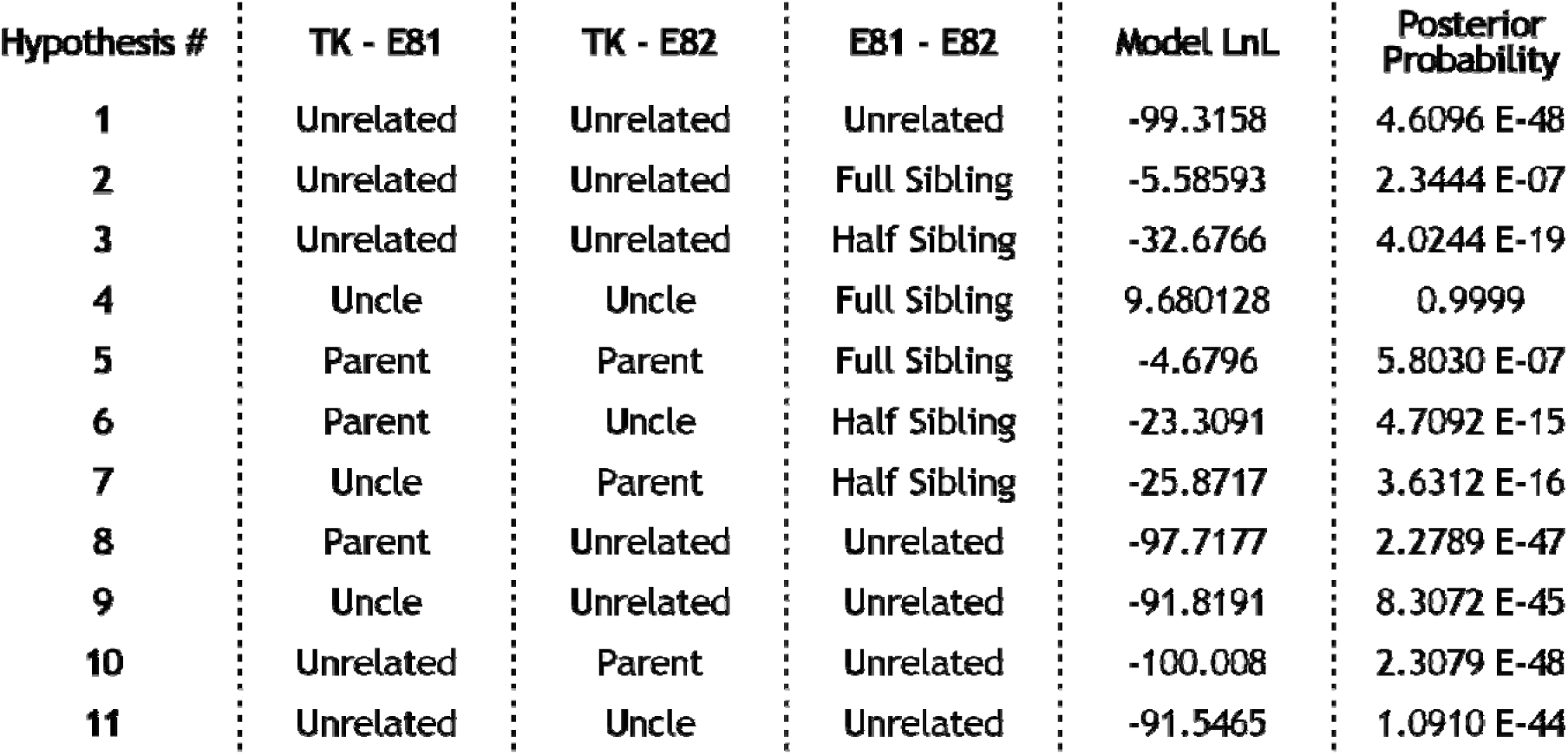
Set of potential relatedness hypotheses for the combinations of full sibling/parental and half sibling/uncle between the three subjects.

| Hypothesis # | TK - E81 | TK - E82 | E81 - E82 | Model LnL | Posterior Probability |
| --- | --- | --- | --- | --- | --- |
| 1 | Unrelated | Unrelated | Unrelated | -99.3158 | 4.6096 E-48 |
| 2 | Unrelated | Unrelated | Full Sibling | -5.58593 | 2.3444 E-07 |
| 3 | Unrelated | Unrelated | Half Sibling | -32.6766 | 4.0244 E-19 |
| 4 | Uncle | Uncle | Full Sibling | 9.680128 | 0.9999 |
| 5 | Parent | Parent | Full Sibling | -4.6796 | 5.8030 E-07 |
| 6 | Parent | Uncle | Half Sibling | -23.3091 | 4.7092 E-15 |
| 7 | Uncle | Parent | Half Sibling | -25.8717 | 3.6312 E-16 |
| 8 | Parent | Unrelated | Unrelated | -97.7177 | 2.2789 E-47 |
| 9 | Uncle | Unrelated | Unrelated | -91.8191 | 8.3072 E-45 |
| 10 | Unrelated | Parent | Unrelated | -100.008 | 2.3079 E-48 |
| 11 | Unrelated | Uncle | Unrelated | -91.5465 | 1.0910 E-44 |

### *In silico* Simulations of Relatedness

In order to assess the accuracy of the relatedness estimations using forced homozygote data, we computed relatedness coefficients using the forced homozygote approach on three relatedness classes – unrelated individuals, first order, and second order, on two different sets of data.

First, we randomly generated SNP data for a total of 2000 virtual pairs of individuals using allele frequencies of the shared SNPs of each of the three possible dyads - TK:E81, TK:E82, E81:E82. For each of these combinations, 2000 unrelated individuals, 2000 full siblings (first order), and 2000 half siblings (second order) were simulated, their relatedness coefficients calculated in SPAGeDI, and the distribution visualised, as shown in Figure 2 (details in the Methods section). The peaks of the curves are at the expected half-values of the relationship coefficients and it is clear that the results obtained for the three relative pairs fall within the expected ranges of variation.

**Figure 2:**
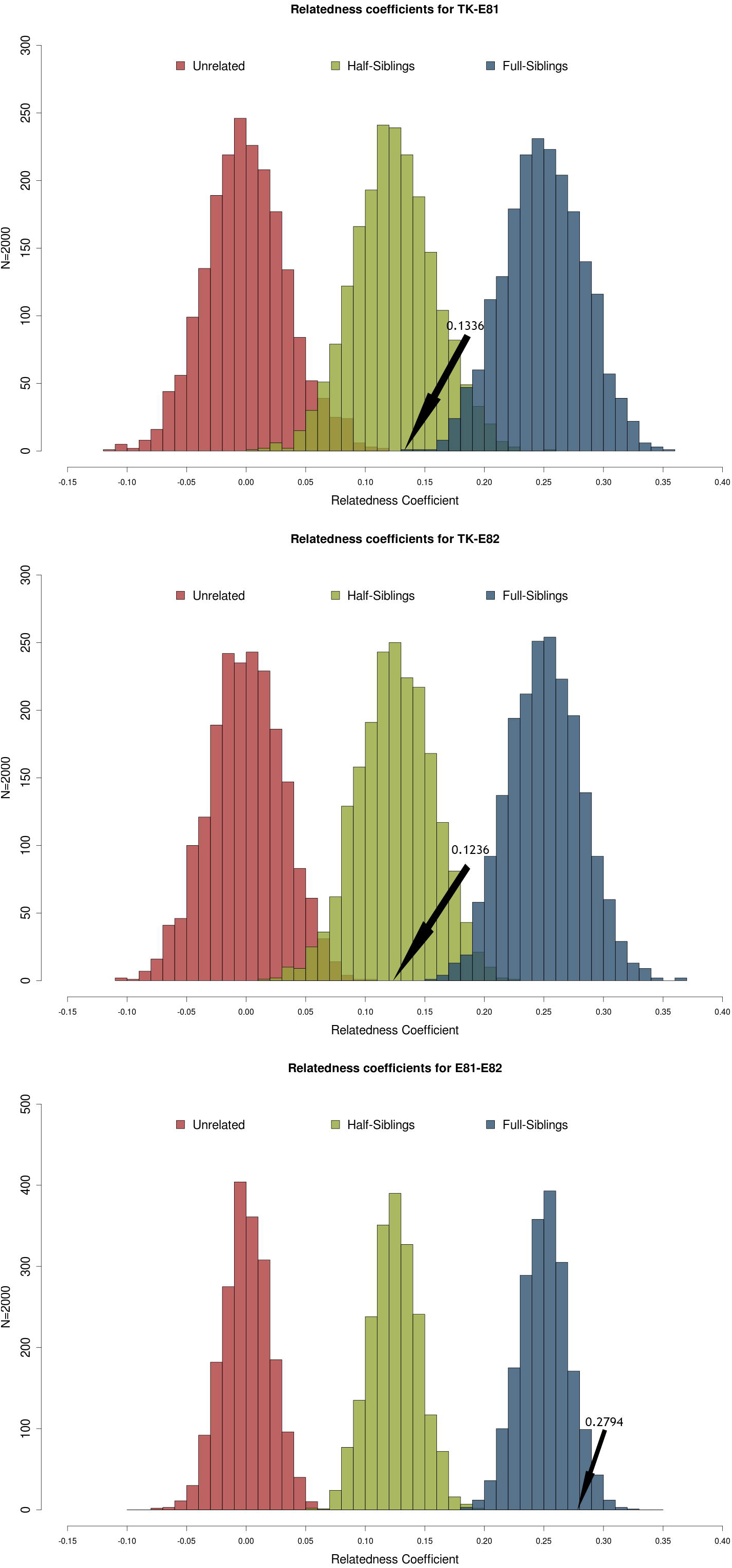
Relatedness coefficients' distribution for Thomas Kent's virtual dyads. “Forced homozygote” relatedness coefficients of computer generated individuals calculated using SPAGeDI1-5a, based on minor allele frequencies of the SNPs common to the pairs TK-E81, TK-E82, E81-E82. Blue-Unrelated, Green-Second Order, Red-First Order. Yellow lines and r values indicate the halved “forced homozygote” relatedness coefficients found for each pair.

We then applied the same approach for three pairs of samples of known relatedness from the 1000 Genomes Project (Table 3), by choosing a pair for each order to test. We randomly downsized each sample to approximately 50.000 SNPs and then ran the simulations for the shared SNPs between each dyad. The number of common SNPs varied from 2040 to 2307, which is in between the values shared by TK and relatives, and one relative and another. The relatedness coefficients for each pair were calculated using exactly the same “forced homozygote” approach and then six hundred estimations per order or relatedness were simulated. These were plotted using the correspondent frequencies of the common SNPs, showing that the coefficients for each pair match their known order of relatedness (Figure 3).

**Figure 3:**
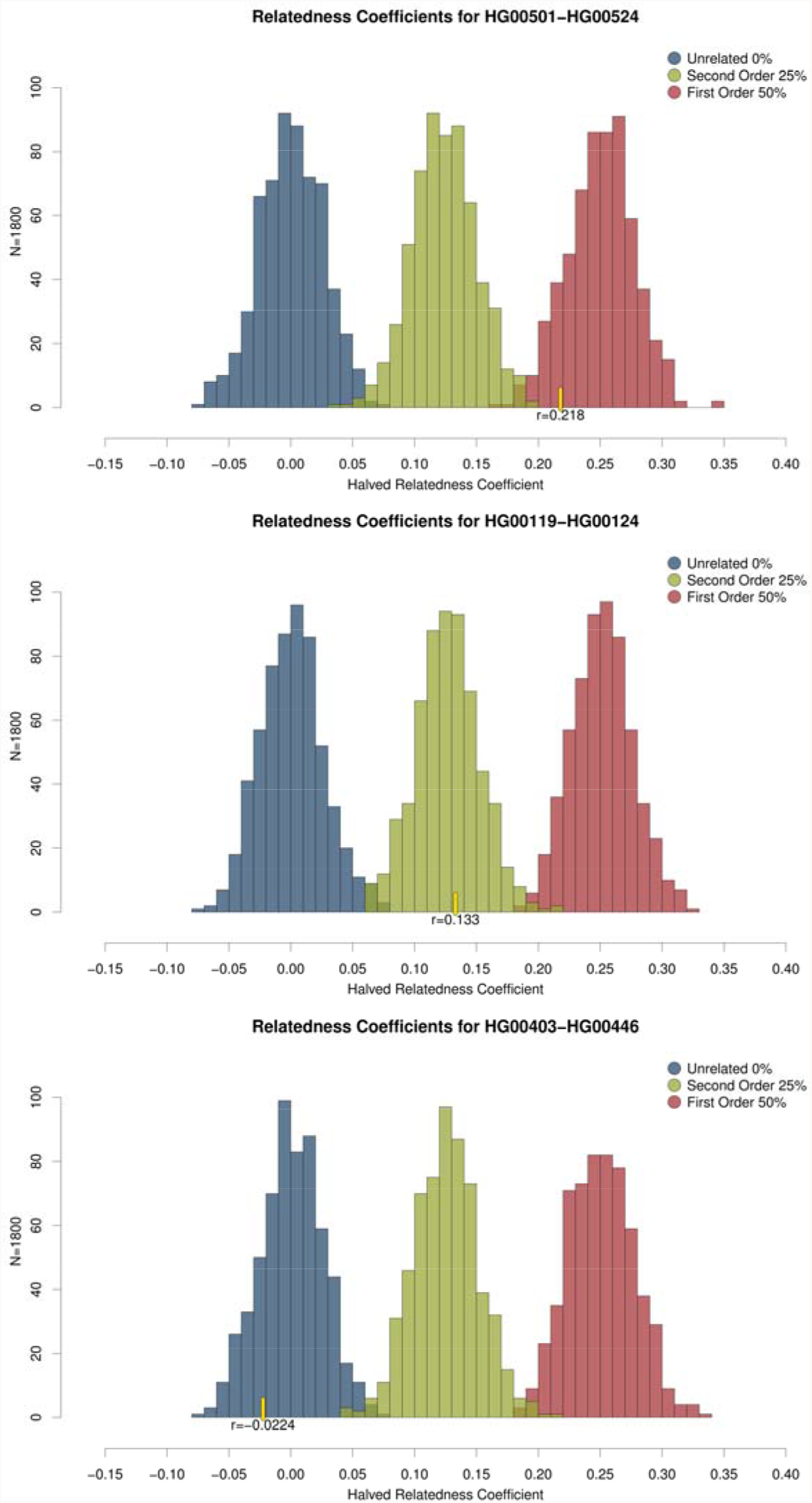
Relatedness coefficients' distribution for the 1000 Genomes Project virtual dyads. “Forced homozygote” relatedness coefficients of computer generated individuals calculated using SPAGeDI1-5a, based on minor allele frequencies of the SNPs common to the pairs HG00501-HG00524, HG00119-HG00124, and HG00403-HG00446. Blue-Unrelated, Green-Second Order, Red-First Order. Yellow lines and r values indicate the halved “forced homozygote” relatedness coefficients found for each pair.

**Table 3:**
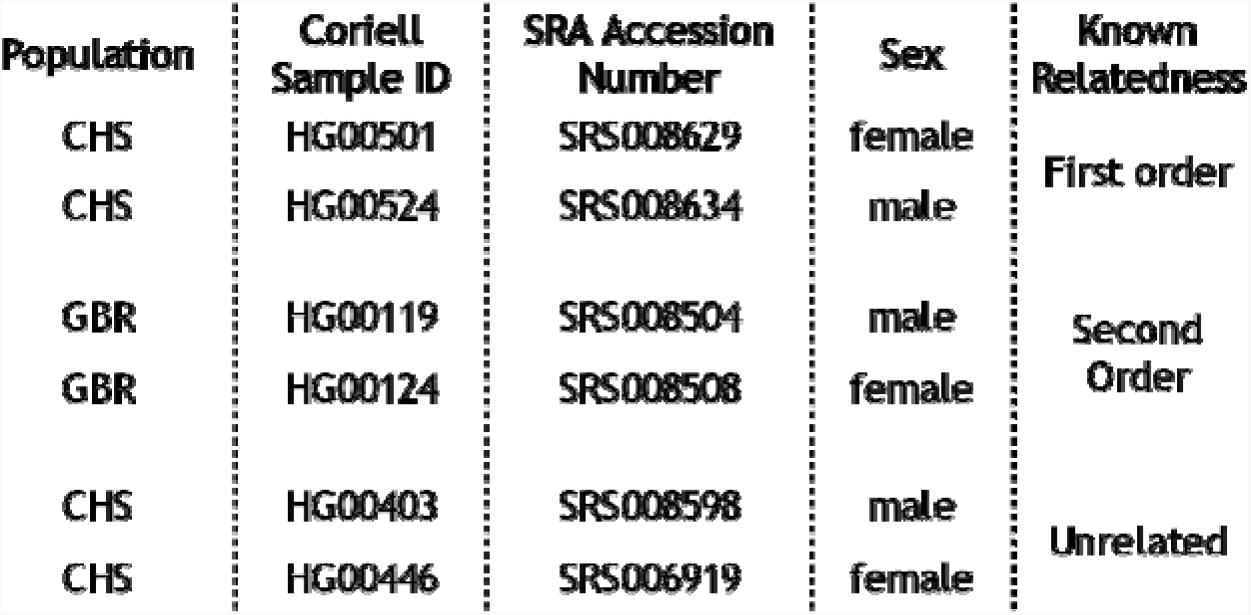
Information on the samples from the 1000 Genomes Project used for testing of the forced homozygote approach.

An R script with two sets of functions (TKrelated and CybRSex) was developed to automate the two processes: simulations down to a true coefficient of relationship of 25%, and actual data tests. By using data in PLINK format as input, our package either runs SPAGeDI for the desired pairs of individuals, or generates X number of homozygous individuals for the given set of SNPs and their allele frequencies. A function allows to plot the three coefficients of relatedness used for simulations (0%, 25%, 50%), making it possible to visualise the distribution of simulated relatedness estimates for any given relatedness class with the expected ranges of variation from the specific input SNP data. The tests on pairs of individuals are performed by a function that requires two input files and an allele frequency file.

This package is freely available under the GNU General Public License v3 at https://github.com/danimag/tkrelated and includes a detailed walkthrough manual.

## CONCLUSIONS

A unique interdisciplinary research opportunity on this historical matter has allowed us to develop an efficient and accurate method for relatedness estimations using small amounts of genetic data. We were able to identify the skeletal remains of Thomas Kent, whose state funeral took place on the 18^th^ of September of 2015, shortly after the identification of his remains. Applicable to both forensic and ancient DNA research, our method for relatedness estimation has important additional benefits in contrast to existing methods. When compared to the software packages PLINK and NGSrelate, the approach we present here requires substantially lower genomic coverage, which will prove helpful when large amounts of genomic data are unavailable, such as in the case of ancient DNA studies. In a situation similar to ours, Korneliussen and Moltke^12^, show that using the software NGSrelate to estimate relatedness based on a coverage of 1X results in large variance of relatedness estimates, yet it still performs better than PLINK. As we have shown, our method was effective with coverages ranging from 0.04X to 0.1X, an order of magnitude reduction on the amount of genetic data required. We also have designed an R script to simulate virtual groups of (un)related individuals and their relatedness coefficients, based on Queller and Goodnights Rxy, from a given set of SNPs and corresponding allele frequencies. This should prove useful in ancient DNA, where low endogenous DNA contents are often the norm and where target enrichment approaches for SNP capture are becoming more common. With the implementation of the “forced homozygote” method, estimating the relatedness between individuals in contexts such as multiple or mass burials may become a more routine task in future studies. This will benefit research in archaeology and anthropology, where the relationships of individuals found interred in multiple burials are often only hypothesized.

## MATERIALS AND METHODS

### Archaeological Bone Sampling

To obtain genetic material from the skeletal remains of Thomas Kent, fine bone powder was retrieved from the cochlea of the left petrous part of the temporal bone that was detached from the rest of the cranium. While the petrous part of the temporal bone is accepted as yielding systematically higher endogenous DNA compared to other skeletal elements^20^, the cochlea in particular was chosen because of research that demonstrated that the otic capsule, and particularly the cochlea, provides the highest endogenous DNA yield from any part of the petrous^14^. The powder was obtained using a minimally-destructive direct drilling technique developed at University College Dublin aimed at reducing any possible damage to the bone. A Dremel 9100 Fortiflex rotary tool, fitted with a small-sized spherical grinding bit (1.5mm) previously treated with bleach and ethanol, was set to medium speed and used to obtain approximately 100mg of bone powder. The cochlea was accessed from the superior aspect of the petrous bone, limiting visible damage to a 2-3mm hole on the superior surface of the petrous. The bone powder generated from drilling the cochlear cavern was collected in a clean weighing boat and transferred to a 1.5mL sterile Eppendorf tube. This procedure was conducted in a clean sample preparation facility at UCD.

### Blood Sampling and DNA Extraction for Modern Relatives

Blood samples were collected from Thomas Kent's living relatives in accordance with the prescribed methods employed by Forensic Science Ireland in the investigation of any unidentified remains. DNA extracts were then sent to University College Dublin for further processing. Informed consent was obtained by the Gardaí for the genetic analysis of this biological material.

### DNA Extraction for Thomas Kent

DNA was extracted from Thomas Kent's bone powder following the protocol from ^22^ which improves upon the optimized silica-based extraction technique described in ^6^. Extraction took place in a physically separated ancient DNA lab at UCD in adherence with stringent anti-contamination protocols. Approximately 50mg of bone powder was combined with 1mL of an extraction buffer solution containing 0.5M EDTA and Proteinase K (Roche Diagnostics). The bone powder was suspended by vortexing and incubated at 37°C with rotation for 18 hours in a ThermoMixer C (Eppendorf AG) and subsequently centrifuged for 2 minutes at 17.000 g in a Heraeus Pico 17 microcentrifuge (Thermo Scientific) to separate the undissolved bone from the supernatant solution. The supernatant solution was collected and added to 13mL of binding buffer solution containing guanidine hydrochloride (MW 95.53, 5M), isopropanol, Tween-20 (10%), and sodium acetate (3M) in a custom-made binding apparatus. This binding apparatus was constructed by forcibly fitting a reservoir removed from a Zymo-Spin V column (Zymo Research) into a MinElute silica spin column (Qiagen). This apparatus was then placed into a 50mL falcon tube^22^. The 14mL solution of binding buffer and DNA extract was added to the extension reservoir in the falcon tube, the cap was secured, and the falcon tube was centrifuged for 4 minutes at 2500rpm, rotated 90°, and centrifuged for another 2 minutes at 3,000rpm. The extension reservoir was then disassembled and the MinElute column was placed into a 2mL collection tube. The column was dry-spun for 1 minute at 13,300rpm, and two wash steps were subsequently performed using 650µL of PE wash buffer. Finally, the column was placed into a clean 1.5mL Eppendorf tube and the DNA was eluted into 25µL of TET buffer.

### DNA Library Preparation

Libraries for next-generation sequencing were built for all three DNA extracts using a modified version of^23^ as outlined in ^20^, where blunt end repair was performed using NEBNext End-Repair (New England Biolabs Inc.) and Bst was inactivated by heat (20 minutes at 80°C). Thomas Kent's DNA library was prepared in a dedicated ancient DNA lab whereas the libraries for the DNA of two modern relatives were prepared in a modern DNA lab in UCD Earth Institute's Area 52. Indexing PCRs were performed with AccuPrime Pfx Supermix (Life Technology), with primer IS4 and an indexing primer. 3µL of the indexed library was added to 21µL of freshly prepared PCR mix, and combined with 1µL of unique index, enabling the pooling of samples for multiplex sequencing. This resulted in a final volume of 25µL. PCR amplification was performed using the following temperature cycling profile: 5 minutes at 95°C, 12 cycles of 15 sec at 95°C, 30 sec at 60°C, and 30 sec at 68°C, and a final period of 5 minutes at 68°C. PCR reactions were then purified using MinElute PCR Purification Kit (Qiagen), following the manufacturer's instructions. Assessment of the PCR reactions were performed on the Agilent 2100 Bioanalyzer following the guidelines of the manufacturer. Based on the concentrations indicated by the Bioanalyzer, samples were pooled in equimolar ratios for sequencing.

### Next-Generation Sequencing

Libraries were sequenced on an Illumina MiSeq platform at the UCD Conway Institute of Biomolecular and Biomedical Research using 65 base pair (bp) single-end sequencing.

### Bioinformatics Analysis

A custom ancient DNA bioinformatics pipeline written by the Pinhasi Lab was applied for processing short length raw MiSeq data. The software cutadapt v1.5^24^ was used to trim adapter sequences. Minimum overlap was set to 1 (- O 1) and minimum length to 17bp (-m 17). Alignment to the human reference genome (hg19, GRCh37) was processed by the Burrows-Wheeler Aligner v.0.7.5a-r405^25^ with disabled seed (-l 1000) and filtering for reads with a minimum phred quality score of 30. Duplicated sequences were removed using samtools v0.1.19-96b5f2294a^26^. To assess the authenticity of Thomas Kent's DNA as ancient, damage patterns were assessed using the mapDamage v.2.0.6 tool^18^.

Single nucleotide polymorphisms were called using the Genome Analyzer Tool Kit's (GATK) v.3.3-0-g37228af Pileup tool for the 354,212 positions present in the Harvard's “Fully public genotype dataset” described in ^10^.

### Relatedness Analysis

Most loci were represented by 1X reads, and this low read depth prevented identification of heterozygote loci for the vast majority of SNP loci in all three analysed individuals, although some loci had greater coverage. These results are the norm in ancient DNA studies, and so we proceeded according to the established protocols. To be able to fully leverage the set of SNP loci, we modeled relatedness on a sample of single-read loci. For loci with greater read depth, we randomly selected one representative allele to reduce the bias that might have been introduced by allowing for some heterozygote loci. By ensuring that all loci contained only one allele we forced a “homozygote” structure on the data. This will necessarily impact the Queller & Goodnight coefficient, as only half the genome is being interrogated, reducing the anticipated relatedness between dyads by a factor of one-half (i.e, reducing first order relationships from 0.5 to 0.25 and second order relationships from 0.25 to 0.125).

#### Thomas Kent Simulations

We reduced the list of genotyped loci to only those loci shared for each dyad (i.e, Thomas Kent and Relative1, Thomas Kent and Relative2, Relative1 and 2). European allele frequencies at the shared loci for each comparison were retrieved from the 1000 genomes project (release 20100804 http://www.1000genomes.org/) using tabix (http://www.htslib.org/doc/tabix.html), and these were used as the reference frequencies for estimating degree of relatedness (symmetrical Rxy estimator, Queller and Goodnight 1989) using SPAGeDi1-5a (build04-03-2015)^21^. This was done using the TKrelated set of functions in the R package developed for this project/approach (detailed walkthrough at https://github.com/danimag/tkrelated). This function reads sample and allele frequencies data in non-binary text PLINK format, *.ped/map, and *.frq, respectively. It then makes that data SPAGeDI-ready, and runs the estimations. It also exports some files that can be used for the virtual simulations.

For the three dyads of relatedness comparison (Thomas Kent and Relative1, Thomas Kent and Relative2, Relative1 and 2), we simulated nine data sets, each with 2000 virtual pairs of full siblings (first order), half siblings (second order) or unrelated individuals, using the observed alleles held in common for the each of the comparisons and the correspondent European allele frequencies (release 20100804 http://www.1000genomes.org/). Within the R package, the set of functions CybRsex take these allele frequencies and generate a desired number of pairs of unrelated individuals, first order relatives, and second order relatives. Each function for each order of relatedness starts by generating random unrelated individuals based on the frequencies of the SNPs from the input file. For first order, it pairs these unrelated individuals and produces one offspring from their homozygous genotypes. The same approach is followed for second order simulations, pairing the common parent with a new unrelated individual. For each simulated data set, we forced the same homozygote condition, resulting in a comparable set of loci represented by one allele. We assessed the degree of relatedness for the simulated data sets with SPAGeDi1-5a. The output relatedness coefficient for each simulated data set was tabulated to create an empirical distribution for three degrees of relatedness (first order, second order, unrelated) for the particular set of loci observed to be held in common for the three subjects.

The distribution of relatedness coefficients was nearly normal (Figure 2). Using mean and variance parameters fit to the empirical distributions, we calculated maximum likelihood (ML) fits of the observed degree of relatedness for each dyad to the three relatedness distributions^27,^^28^. Potential relatedness hypotheses constitute the set of potential combinations of full sibling/parental and half sibling/uncle between the three subjects, producing a set of eleven potential hypotheses (Table 2). The ML fit of each hypothesis is then the sum of the ML fits of the observed relatedness coefficient between the two individuals for the appropriate empirical distribution.

#### 1000 Genomes Simulations

To test the robustness of our approach, we applied it to three pairs of individuals with know relatedness from the 1000 Genomes Project. Since related individuals are excluded in Phase 3, we downloaded the variant calls from Phase 1 for all chromosomes (release 20101123 from http://www.internationalgenome.org/data, accessed on 27/09/2016). Using PLINK v.1.90b3.41^11^ we converted and merged the data. We selected the individuals shown in Table 3. They were isolated from the dataset, randomly sub-sampled to around 50.000 SNPs, and we then ran our script for estimating relatedness and simulating individuals. As our approach has been designed to be used with samples from very low-coverage scenarios such as in ancient DNA studies, we ran these tests with approximately 2000 SNPs and 600 simulations. We retrieved allele frequencies from the populations from where each pair of individuals was originated, i.e. for the second order test on a pair of individuals originating from Great Britain we used the allele frequencies of that same population. The results of these simulations confirm the robustness of our approach when dealing with low-coverage data (Figure 3).

### ETHICS STATEMENT

The investigation into the authentication of Thomas Kent's remains was tasked to the Irish Police, An Garda Siochana, on behalf of the State, and therefore obliged to adhere to specific ethical and legal considerations.

Informed consent was obtained when collecting the blood from Thomas Kent's living relatives in regard to analysis of the genetic data and dissemination of the results. Kent's remains, legally considered archaeological, were handled with the permission of the correspondent legal authorities

The request for assistance from UCD by An Garda Siochana to identify the remains recovered from Cork Prison in early 2015 was made to progress that element of the overall investigation.

An Garda Siochana are tasked with such investigations, on behalf of the State, and do not require an ethics committee to initiate enquiries. An Garda Siochana may enlist the expertise of any agency or academic entity to pursue lines of enquiry, and such was the case with UCD.

In this case, the original request for help in identifying the remains came from the Department of An Taoiseach (Head of Government) to the National Forensic Coordination Office, who then managed the overall investigation. The integrity of all evidence, samples and results was managed by the Head of the National Forensic Coordination Office, who was also the investigating officer in this case. All ethical considerations and legal obligations under the Data Protection Acts were his responsibility as Investigating Officer, and he was the person who sought the assistance of UCD on behalf of An Garda

Siochana. He reviewed all evidence or results before they were communicated to the relevant parties. In such investigations ethical considerations form part of the overall review, in addition to many layers of legal consideration and all requirements were met.

## DATA AVAILABILITY

The genetic data from the study has been stored in the repository of the National Forensic Coordination Office of An Garda Siochana, and although it is of public access, legal considerations require that it complies with the Data Protection Act, making it restricted. The data will be available from the Police repository by request, under the reference number NFCO-01-244103/15. The following email can be used.

## ACKNOWLEDGEMENTS

We would like to thank Dr. Sudipto Das for his comments on sequencing methods; Dr. Eppie Jones for comments on the manuscript; the Irish Government for their support throughout the Thomas Kent identification process; and Dr. Olivia Cheronet for helping with the R script. This research was supported by R.P.'s European Research Council Starting grant ERC- 2010-StG 263441 (https://erc.europa.eu). D.F. was supported by an Irish Research Council Post-Graduate grant GOIPG/2013/36 (www.research.ie).

## AUTHOR CONTRIBUTIONS

D.F., J.C, K.S., M.N. and R.P. designed the experiments. D.F., E.C., J.E.C, K.S. and M.N. carried the experimental work. D.F., E.F., J.C., J.F. and K.S. analysed the data. D.F., J.B., J.C., J.F and K.S. wrote the manuscript. Tables and Figures were created by D.F.

## COMPETING FINANCIAL INTERESTS

The authors declared no competing financial interests.

## References

1. Powell, J. E., Visscher, P. M. & Goddard, M. E. Reconciling the analysis of IBD and IBS in complex trait studies. Nat Rev Genet 11, 800–805 (2010).

2. Speed, D. & Balding, D. J. Relatedness in the post-genomic era: is it still useful? Nat Rev Genet 16, 33–44(2015).

3. Pääbo, S. Ancient DNA: extraction, characterization, molecular cloning, and enzymatic amplification. Proceedings of the National Academy of Sciences 86, 1939–1943(1989).

4. Mitchell, D., Willerslev, E. & Hansen, A. Damage and repair of ancient DNA. Mutation Research/Fundamental and Molecular Mechanisms of Mutagenesis 571, 265–276(2005).

5. Willerslev, E. et al. Long-term persistence of bacterial DNA. Current Biology 14, R9–R10 (2004).

6. Rohland, N. & Hofreiter, M. Ancient DNA extraction from bones and teeth. Nat. Protocols 2, 1756–1762(2007).

7. Baca, M., Doan, K., Sobczyk, M., Stankovic, A. & Węgleński, P. Ancient DNA reveals kinship burial patterns of a pre-Columbian Andean community. BMC Genetics 13, 1–11(2012).

8. Deguilloux, M. F. et al. Ancient DNA and kinship analysis of human remains deposited in Merovingian necropolis sarcophagi (Jau Dignac et Loirac, France, 7th–8th century AD). Journal of Archaeological Science 41, 399–405(2014).

9. Dudar, J. C., Waye, J. S. & Saunders, S. R. Determination of a kinship system using ancient DNA, mortuary practice, and historic records in an upper Canadian pioneer cemetery. International Journal of Osteoarchaeology 13, 232–246(2003).

10. Haak, W. et al. Massive migration from the steppe was a source for Indo-European languages in Europe. Nature 522, 207–211(2015).

11. Chang, C. C. et al. Second-generation PLINK: rising to the challenge of larger and richer datasets. GigaScience 4, 1–16(2015).

12. Korneliussen, T. S. & Moltke, I. NgsRelate: a software tool for estimating pairwise relatedness from next-generation sequencing data. Bioinformatics 31, 4009–4011(2015).

13. Barton, B. The Secret Court Martial Records of the Easter Rising. (The History Press Ireland, 2010).

14. Pinhasi, R. et al. Optimal Ancient DNA Yields from the Inner Ear Part of the Human Petrous Bone. PloS one 10, e0129102 (2015).

15. Mathieson, I. et al. Genome-wide patterns of selection in 230 ancient Eurasians. Nature 528, 499–503(2015).

16. Queller, D. C. & Goodnight, K. F. Estimating relatedness using genetic markers. Evolution 43, 258–275(1989).

17. Ginolhac, A., Rasmussen, M., Gilbert, M. T. P., Willerslev, E. & Orlando, L. mapDamage: testing for damage patterns in ancient DNA sequences. Bioinformatics 27, 2153–2155(2011).

18. Jónsson, H., Ginolhac, A., Schubert, M., Johnson, P. L. F. & Orlando, L. mapDamage2.0: fast approximate Bayesian estimates of ancient DNA damage parameters. Bioinformatics 29, 1682–1684(2013).

19. Navarro-Gomez, D. et al. Phy-Mer: a novel alignment-free and reference-independent mitochondrial haplogroup classifier. Bioinformatics 31, 1310–1312(2015).

20. Gamba, C. et al. Genome flux and stasis in a five millennium transect of European prehistory. Nature communications 5, 5257 (2014).

21. Hardy, O. J. & Vekemans, X. spagedi: a versatile computer program to analyse spatial genetic structure at the individual or population levels. Molecular Ecology Notes 2, 618–620(2002).

22. Dabney, J. et al. Complete mitochondrial genome sequence of a Middle Pleistocene cave bear reconstructed from ultrashort DNA fragments. Proceedings of the National Academy of Sciences 110, 15758–15763(2013).

23. Meyer, M. & Kircher, M. Illumina sequencing library preparation for highly multiplexed target capture and sequencing. Cold Spring Harbor protocols 2010, pdb prot5448 (2010).

24. Martin, M. Cutadapt removes adapter sequences from high-throughput sequencing reads. 2011 17, 10–12(2011).

25. Li, H. & Durbin, R. Fast and accurate short read alignment with Burrows–Wheeler transform. Bioinformatics 25, 1754–1760(2009).

26. Li, H. et al. The Sequence Alignment/Map format and SAMtools. Bioinformatics 25, 2078–2079(2009).

27. Edwards, A. W. F. Likelihood: Expanded Edition. (The Johns Hopkins University Press, 1992).

28. Royall, R. M. Statistical Evidence: A Likelihood Paradigm. (Chapman and Hall, 1997).

